# Modeling non-genetic dynamics of cancer cell states measured by single-cell analysis: Uncovering bifurcations that explain why treatment either kills a cancer cell or makes it resistant

**DOI:** 10.1101/574848

**Authors:** S. Huang

## Abstract

Single-cell transcriptomics offers a new vista on non-genetic tumor cell plasticity, a neglected aspect of cancer. The gene expression state of each cell is governed by the gene regulatory network which represents a high-dimensional non-linear dynamical system that generates multiple stable attractor states and undergoes destabilizing bifurcations, manifest as critical transitions. Modeling clonal cell population as statistical ensembles of the same dynamical system, a index *IC* is derived for detecting destabilization towards critical transitions in single-cell molecular profiles. Therapy-induced bifurcation explains why treatment backfires: a drug-treated cell is imposed the binary choice to either apoptose or become a cancer-stem cell.

## I. Introduction to the type of problem in cancer

Tumor cell populations within a tumor exhibit a vast heterogeneity of cell states and rapid bidirectional interconversions; a behaviors is not readily explained by genetic mutations. Such non-genetic plasticity allows cancer cells to spontaneously or in response to environmental cues switch between discretely distinct stable phenotypes, such as a cancer stem cell (CSC), proliferative, differentiated, senescent or apoptotic states [1]. Such cell state dynamics allows cells to switch from a drug-sensitive to a drug-resistant state. We model the ubiquitous unintended drug-induced conversion of cells that survive treatment into a drug-resistant, CSC-like state as an inevitable consequence of bifurcation in a dynamical system [2].

### Epistemological note on mathematical oncology

Sadly, most cancer research promoted by current funding for interdisciplinary approaches uses mathematical tools to address not scientific but operational problems (e.g. optimization of therapy schedules, image analysis) and typically employ *heuristics* (*ad hoc* models “to see if it works”). Such mathematical modeling is designed in the mind-set of engineering, not of basic science. While sometimes useful for making practical predictions, such heuristics do not seek to *explain* observable phenomena in terms of first principles of natural sciences. In contrast to engineering, science aspires to explain biological observables, such as interesting, reoccurring (“universal”) tumor behaviors, by a formal hypothesis and followed by demonstration, that they are the *necessary* manifestation of constraints imposed by underlying principles of biochemistry, molecular biology, cell population dynamics, etc. Such formal “bottom-up” approaches of course will still require coarse-graining in the formalization of a model that however is rooted in a theory of natural science and must not be confounded with the above heuristic, *ad hoc* models.

## II. Results of application of Method

### A. The model class: multi-stable dynamical system

One such coarse-graining that formalizes the constraints in terms of governing principles is the use of dynamical systems theory to describe changes in gene expression profiles which determine the cell phenotype, defined by a set of *m* genes *i: **x**(t*) = [*x*_1,_ *x*_2.._*x*_*i . .*_*x*_*m*_]. The dynamical system captures the constraints unto the time evolution of *x (t*) by a (genomically encoded) gene regulatory network (GRN) through which these *m* genes interact. The state vector **x** represents a state of the GRN at time *t*, thus one point in the *m*-dimensional state space. We have then the *m*-dimensional dynamical system 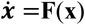. Herein, functions *F*_*i*_ captures at a high-level the GRN architecture and how each the expression behavior of each gene *i* is influenced by the values *x*_*j*_ of its “upstream” regulators. The system is multi-stable: it exhibits multiple stable attractor states 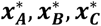 … One (simplified) premise for which there is now ample experimental support [3, 4], is that observable, robust (recurring, characteristic) expression profiles, such as those defining the normal cell types in the body or the distinct cancer cell states within a tumor, correspond to these high-dimensional stable attractor states of 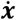. Gene expression noise or perturbations can then cause switch-like transitions between attractors (phenotype conversions) [4].

We depart from standard dynamical systems modeling efforts in two ways: (*i*) Since the functional form of **F** is generally not known we do not explicit model ***x**(t*) but focus on generic observable features associated with a *critical state transitions* [5]. This leads to a “phenomenological” model that is agnostic of specifics of the GRN and cannot predict the trajectory **x**(*t*) but predicts generic properties that can be experimentally tested. (*ii*) We take advantage of single-cell analysis technology to measure the expression of gene *i* in cell *k*: *x*_*ik*_. Thus, we obtain the expression state **x**_*j*_ in each individual cell *j* of a population of *n* cells which are nominally identical (clonal *and* in the same phenotypic state **x**) –distinct only because of gene expression noise. We then assume an ergodic ensemble of *n* replicates of our dynamical system: ***x***_1_(*t*), ***x***_2_(*t*),.. ***x***_*n*_(*t*). For each dimension *i* the value ***x***_*i*_ is subject to stochastic fluctuations, such that gene *i* across the cell population represents a distribution *P*_*i*_(*x*_*i*_). Thus, at any time point we have a *cell population state*, and instead of the state vector **x**(*t*) we have the matrix **X**(*T*) with elements ***x***_*ik*_ which can be approximately measured with single-cell qPCR or RNAseq as population snapshot of time *T*:

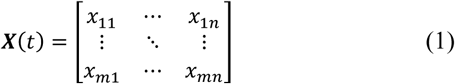

### B. The formal hypothesis

We model the scenario in which a perturbation on **X** (treatment) to push cells into the apoptosis attractor also produces stem-cell states in those cells not killed.

The *central hypothesis* is that bifurcations drive attractor switching, akin to catalysis lowering “energy barriers” (Fig.1). Cell phenotype switching is a response to an environmental signal that operates an unknown bifurcation parameter μ whose change destabilizes the current attractor 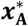, hence opening access to 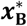. Thus, the attractor transition represents a *critical state transition* [5]. If the bifurcation is of a pitch-fork-type (a sufficient but not necessary condition in a multi-stable, rugged “energy” landscape [6]), then upon destabilization of 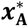, not necessarily only the desired attractor state is accessed by the unstable cells, but due to stochastic fluctuations, other nearby attractors (the alternative branch in a pitch-fork-type bifurcation) could also be accessed. These undesired cells would explain “rebellious cells” that switch to states opposite to the intended one [2]; in the case of cancer treatment as the bifurcation inducer, 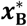 would correspond to the desired destination state (apoptosis) where as the “opposite” state 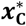 would represent the treatment induced stem-like state. This would explain why cancer therapy often backfires–generating cancer stem cells [1].

**Figure 1.**
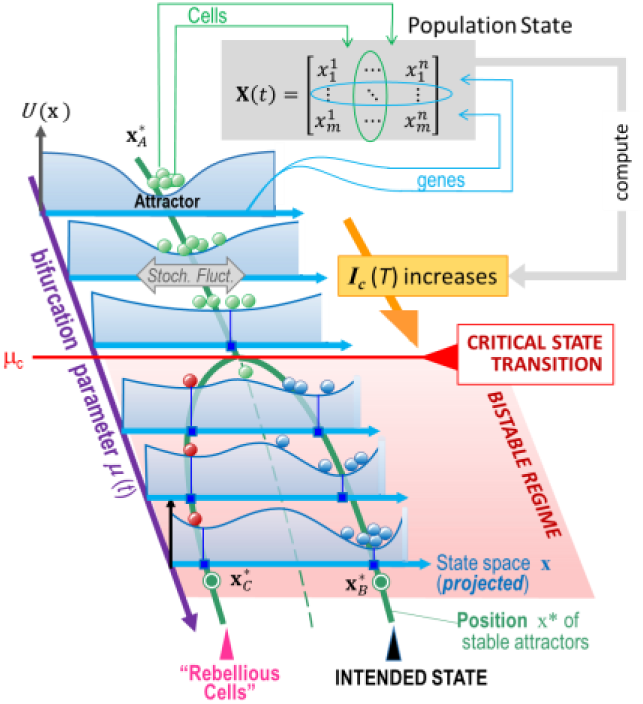
Schematic of model and hypothesis based on a pitchfork bifurcation (as pedagogically simple example). For landscape “elevation”, *U*(*x*) see [6]

### C. Application of the model

The goal of applying this model to experimental data is to test the key assumption: the presence of a destabilization prior to the induced transition to the new attractor. Without knowledge of **F** the phenomenological approach is to predict changes in **X**(*T*) associated with approach to a critical state transitions, analogous to the Early Warning Signals for critical transitions in low-dimensional systems [5] monitored continuously in time, such as the enhancement of fluctuations (auto-correlation in time). Here the equivalent, under the assumption of ergodicity, would be a particular change in the structure of high-dimensional population structure captured in **X**(*T*). We derived the “critical transition index” *I*_*C*_ [2] which has the key property that *I*_*C*_ will increase monotonically if a system (monostable cell population) moves towards a critical transition:

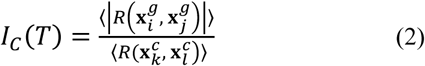

where 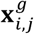 are gene vectors (rows in eq. (1)), and 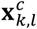 are cell state vectors (columns) and <…> denotes the average of Pearson coefficients *R* for all pairs of genes (*i,j*) or cells (*k, l*). That the value of the denominator will decrease when a critical transition is imminent is obviously a consequence of the increased diversity of cell states as stability is reduced, reflecting the “critical slowing down” [5] of relaxation to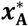. The less obvious increase of the numerator is related to the reduced dimensionality of the manifold in the *m*-dimensional state space at bifurcation –see Appendix in [2]. Experimental verification by single-cell qPCR measurements showing increase in *I*_*C*_ in a case of known cell fate bifurcation is presented in ref. [2].

## III. Quick Guide to the Method

### A. Model assumptions

To derive *I*_*C*_ we made the following basic assumptions [2]: (*i*) Any robust biologically functional cell state represents a hyperbolic stable attractor state that is attracting in all *m* (most?) dimensions of GRN dynamics. (*ii*) Ergodicity and quasisteady state process: expression levels *x*_*i*_ is subject to stochastic fluctuations that are much faster than the gradual change in the bifurcation parameter μ. As a consequence of both assumptions, the cell population to be measured to compute *I*_*C*_ should be a mono-modal cell population.

### B. Application to single-cell expression analysis

The presented method of computing *I*_*C*_ (2) from the single-cell profiling matrices **X**(*T*) (1) for a population of *n* cells with their states described by *m* genes can be applied to the analysis of single-cell resolution any molecular profiles in (tumor) cell populations, obtained by new technologies, such as single-cell transcriptomics (RNAseq and qPCR), CyTOF [7] and *in situ* methods. Few time points *T* suffice to detect a trend of *I*_*C*_ indicating destabilization towards a critical transition which may be new indicator for cancer progression.

## Notes

Research supported by the NIH (NIGMS) and the Canadian NSERC and CIHR. S. Huang is at the Institute for Systems Biology, Seattle WA 98109, USA.

## References

[1] Pisco, A. O. & Huang, S. “Non-genetic cancer cell plasticity and therapy-induced stemness in tumour relapse: ‘What does not kill me strengthens me’”. Br J Cancer 112, pp. 1725–1732 (2015)

[2] Mojtahedi, M. et al. “Cell Fate Decision as High-Dimensional Critical State Transition”. PLoS Biol 14, e2000640 (2016)..

[3] Chang, H. H., et al. “Transcriptome-wide noise controls lineage choice in mammalian progenitor cells”. Nature 453, 544–547 (2008.

[4] Huang, S. “Systems biology of stem cells: …”. Philos Trans R Soc Lond B Biol Sci 366, pp. 2247–2259 (2011).

[5] Scheffer, M. et al. “Early-warning signals for critical transitions”. Nature 461, pp. 53–59 (2009).

[6] Zhou, J. X., et al. “Quasi-potential landscape in complex multi-stable systems". J Royal Soc Interface 9, pp 3539–3553 (2012).

[7] Marr C, Zhou JX, Huang S. “Single-cell gene expression profiling and cell state dynamics: collecting data, correlating data points and connecting the dots”. Curr Opin Biotechnol. 39 pp. 207–214 (2016)

